# CREAM: Clustering of genomic REgions Analysis Method

**DOI:** 10.1101/222562

**Authors:** Seyed Ali Madani Tonekaboni, Parisa Mazrooei, Victor Kofia, Benjamin Haibe-Kains, Mathieu Lupien

**Author notes:** Corresponding author: Seyed Ali Madani Tonekaboni; Benjamin Haibe-Kains; Mathieu Lupien.

## Abstract

Cellular identity relies on cell type-specific gene expression profiles controlled by cis-regulatory elements (CREs), such as promoters, enhancers and anchors of chromatin interactions. CREs are unevenly distributed across the genome, giving rise to distinct subsets such as individual CREs and Clusters Of cis-Regulatory Elements (COREs), also known as super-enhancers. Identifying COREs is a challenge due to technical and biological features that entail variability in the distribution of distances between CREs within a given dataset. To address this issue, we developed a new unsupervised machine learning approach termed Clustering of genomic REgions Analysis Method (CREAM) that outperforms the Ranking Of Super Enhancer (ROSE) approach. Specifically CREAM identified COREs are enriched in CREs strongly bound by master transcription factors according to ChIP-seq signal intensity, are proximal to highly expressed genes, are preferentially found near genes essential for cell growth and are more predictive of cell identity. Moreover, we show that CREAM enables subtyping primary prostate tumor samples according to their CORE distribution across the genome. We further show that COREs are enriched compared to individual CREs at TAD boundaries and these are preferentially bound by CTCF and factors of the cohesin complex (e.g.: RAD21 and SMC3). Finally, using CREAM against transcription factor ChIP-seq reveals CTCF and cohesin-specific COREs preferentially at TAD boundaries compared to intra-TADs. CREAM is available as an open source R package (https://CRAN.R-project.org/package=CREAM) to identify COREs from cis-regulatory annotation datasets from any biological samples.

## BACKGROUND

Over 98% of the human genome consists of sequences lying outside of gene coding regions that harbor functional features, including cis-regulatory elements (CREs), important in defining cellular identity [1, 2]. CREs such as enhancers, promoters and anchors of chromatin interactions, are predicted to cover 20-40% of the noncoding genomic landscape [3]. CREs define cell type identity by establishing lineage-specific gene expression profiles [4–6]. Current methods to annotate CREs in any given biological sample include ChIP-seq for histone modifications (e.g., H3K4me1, H3K4me3 and H3K27ac) [4,5,7], chromatin binding protein (e.g., MED1, P300, CTCF and ZNF143) [7–9] or through chromatin accessibility assays (e.g., DNase-seq and ATAC-seq) [10, 11].

Clusters Of cis-Regulatory Elements (COREs), such as clusters of open regulatory elements, super-enhancer and stretch-enhancers, were recently introduced as a subset of CREs based on different parameters including close proximity between individual CREs [12–15]. COREs are significantly associated to cell identity and are bound with higher intensity by transcription factors than individual CREs [13,14,16]. Furthermore, inherited risk-associated loci preferentially map to COREs from disease related cell types [12,17–19]. Finally, COREs found in cancer cells lie proximal to oncogenic driver genes [20–23]. Together, these features showcase the utility of classifying CREs into COREs versus individual CREs.

Recent work assessed the role of COREs as a collection of individual CREs proximal to each other as opposed to a community of synergizing CREs [24–26]. Partial redundancy between effect of individual CREs versus a CORE on regulating expression of genes in embryonic stem cells was observed [24] as well as low synergy between the individual CREs within COREs [27]. Whether COREs provide an added value over individual CREs to gene expression is still debated. Conclusions may be confounded by the simplistic approach commonly used to identify COREs. For instance, the distance between CREs is a critical feature that distinguishes COREs from individual CREs. Available methods to identify COREs dismiss the variability in the distribution of distances between CREs that stems from technical and biological features unique to each CRE dataset. Instead, arbitrary thresholds are considered including 1) a fixed stitching distance limit between CREs (such as 20 [15] or 12.5 [13] kilobases) to report them within a CORE, 2) a fixed cutoff in the ChIP-seq signal intensity from the assay used to identify CREs to separate COREs from individual CREs [13], or 3) reporting an individual CRE with high signal intensity as a CORE [13, 28]. To address these limitations, we developed a new methodology termed CREAM (Clustering of genomic REgions Analysis Method) (Fig. 1). CREAM is an unsupervised machine learning approach that takes into account the distribution of distances between CREs in a given biological sample to identify COREs, consisting of at least two individual CREs.

**Figure 1.**
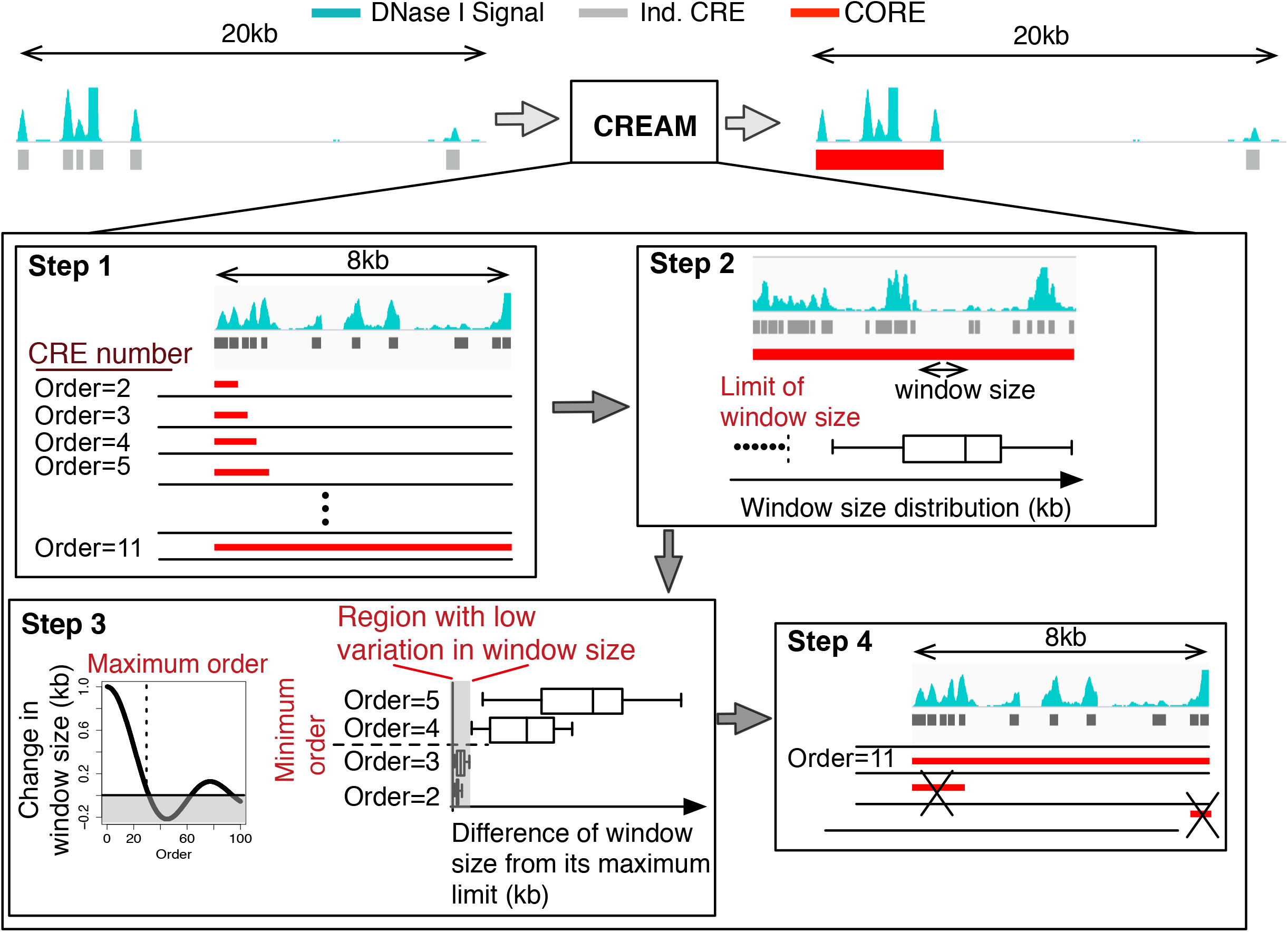
Schematic representation of the four main steps of Clustering of genomic REgions Analysis Method (CREAM): Step 1) CREAM identifies all clusters of 2, 3, 4 and more neighboring CREs. The total number of CREs in a cluster defines its “Order”; Step 2) Identification of the maximum window size (MWS) between two neighboring CREs in clusters for each Order. The MWS corresponds to the greatest distance between two neighboring CREs in a given cluster; Step 3) identification of maximum and minimum Order limits of COREs from a given dataset; Step 4) CORE reporting according to the criteria set in step 3 from the highest to the lowest Order.

Benchmarking CREAM against Rank Ordering of Super-Enhancers (ROSE) [20, 29], the current standard method to call COREs, we demonstrate, using DNase-seq data, that CREAM outperforms ROSE to identify COREs proximal to highly expressed genes, associated with high intensity transcription factor binding, predictive of cell identity and able to subtype primary prostate tumor samples. We further show that CREAM identified COREs enrich proximal to genes essential for the growth of cancer cells. Finally, by integrating maps of the three-dimensional architecture of the human genome with COREs, we reveal the enrichment of CTCF and cohesin COREs found in Topologically Associated Domain (TAD) boundaries.

## RESULTS

ENCODE generated DNAse-seq data from GM12878 and K562 cell lines were used to characterize the COREs identified by CREAM and ROSE. We focused on these cell lines because of their extensive characterization by the ENCODE project [30], inclusive of expression profiles and DNA-protein interactions assessed by ChIP-seq assays, allowing for a comprehensive biological assessment of COREs identified in each cell line. CREAM identified a total of 1,694 and 4,968 COREs from DNAse-seq in GM12878 and K562 cell lines, respectively. These COREs account for 14.6% and 17.2% of all CREs reported by the DNase-seq profiles from these cells, respectively. ROSE identified 2,490 and 2,527 COREs in GM12878 and K562, respectively, accounting for 31% and 30% of CREs found in each cell line. We next assessed the exclusivity of COREs identified by CREAM and ROSE in GM12878 or K562 cell lines. Approximately 85% of CREAM-identified COREs in GM12878 and 49% in K562 overlap with ROSE-identified COREs (Fig. 2A). Conversely, only 14% of ROSE identified COREs in GM12878 cells and 8% in K562 cells overlap CREAM-identified COREs (Fig. 2A). In addition, COREs identified by ROSE tend to be larger (average 41kb and 138 kb width in GM12878 and K562, respectively) than those reported by CREAM (average 9kb and 5kb width in GM12878 and K562, respectively) (Fig. 2B). Hence, the CREAM and ROSE methods differ sufficiently to delineate distinct types of COREs warranting further comparison.

**Figure 2.**
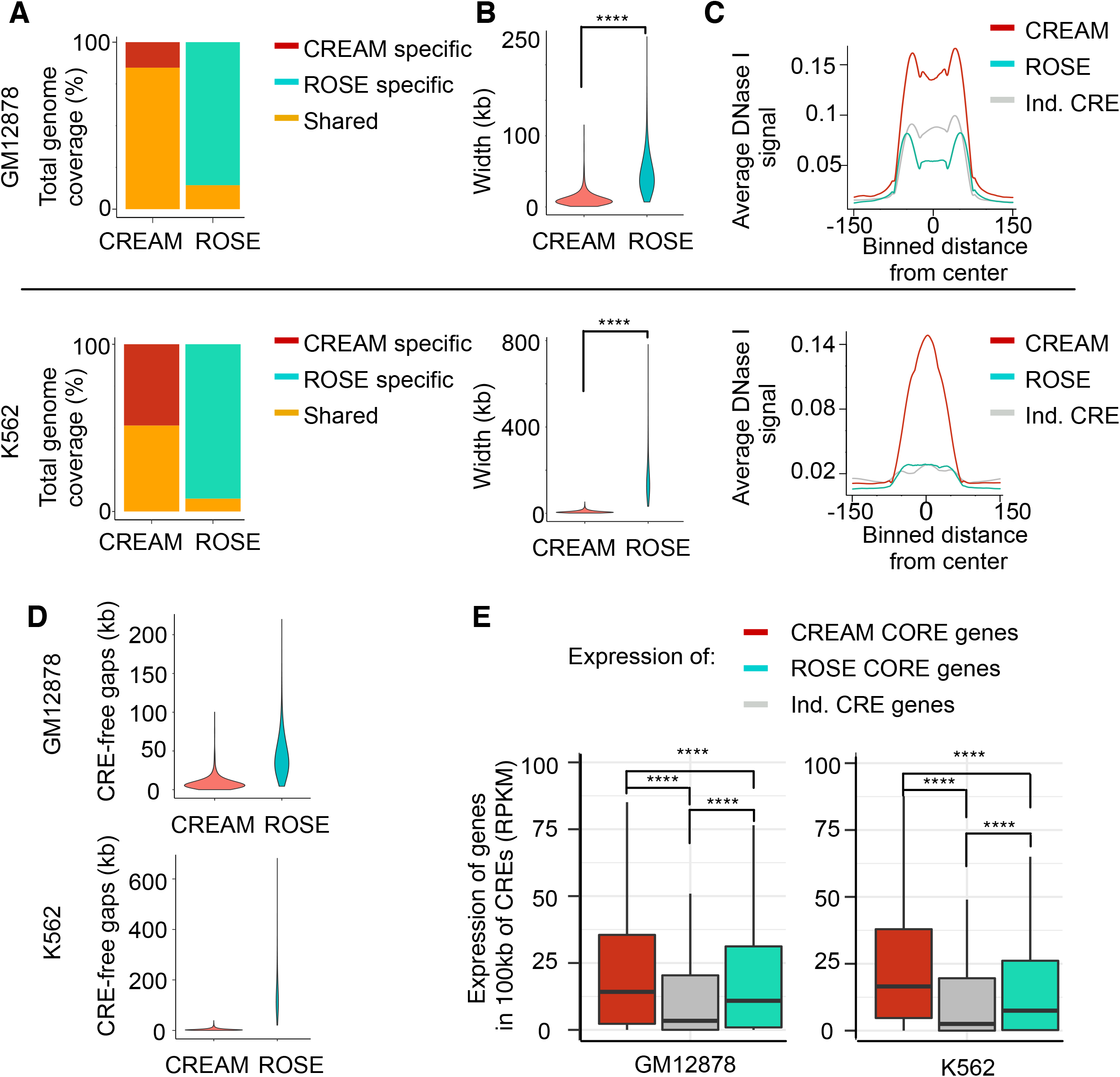
Comparison of genomic characteristics of the COREs identified by CREAM versus ROSE in GM12878 and K562 cell lines. **(A)** Percentage of the identified COREs by CREAM and ROSE which are unique or shared regarding the genomic coverage. (**B**) Distribution of CORE widths. (**C**) Enrichment of DNase I signal profile in individual CREs and COREs. Each CORE (or individual CREs) plus its flanking regions are binned into 100 bins each. (**D**) Size of CRE-free gaps within COREs. (**E**) Expression level of genes associated to individual CREs or COREs.

### DNase I hypersensitive signal is elevated within COREs

COREs are reported to associate with higher levels of binding for a wide range of chromatin binding proteins [14]. Our results show that COREs identified by CREAM have 2 to 5 fold higher average DNase I hypersensitivity signal per base pair (bp) compared to individual CREs in either cell line (Fig. 2C). This is in contrast to COREs identified by ROSE, which show an equivalent DNase I hypersensitivity to individual CREs in K562 cells and less than a 1.5 fold increased in GM12878 cells (Fig. 2C). The distinct behavior between CREAM and ROSE-identified COREs could be due to their size difference that translates in more base pairs free of CRE (CRE-free gaps) in ROSE-identified COREs (102 mbp and 208 mbp in GM12878 and K562 cells, respectively) compared to CREAM-identified COREs (14.7 mbp and 18.7 mbp in GM12878 and K562 cells, respectively) (Fig. 2D). This stems from a permissive <12.5kb distance between CREs criteria in ROSE. In contrast, a learned maximum distance limit between CREs criteria is used by CREAM resulting in smaller average of maximum distances (<1.7 kb in GM12878 and <1 kb in K562 cells for their respective DNase-seq delineated CREs).

### CREAM-identified COREs are proximal to highly expressed genes

In agreement with previous reports [13, 14], COREs identified by ROSE are proximal to genes expressed at higher levels than those near individual CREs (Fig. 2E). This also applies to COREs identified by CREAM in both GM12878 (>4 fold difference) and K562 cell lines (>2.5 fold difference) (Fig. 2E). Noteworthy, genes proximal to CREAM-identified COREs have at least a 1.5 fold higher expression levels compared to genes proximal to ROSE-called COREs (p<0.001) in either cell line (Fig. 2E). We further assessed expression of genes in proximity of COREs specific to CREAM and ROSE. Expression of genes in proximity of CREAM-specific COREs were significantly higher than genes in proximity of ROSE-specific COREs in both GM12878 and K562 cell lines (p<0.001) (Supplementary Fig. 1). Moreover, comparing the percentage of COREs which overlap with promoters, exons, introns, and intergenic regions reveals a very similar distributions for COREs identified by CREAM or ROSE in GM12878 and K562 cell lines. (Supplementary Fig. 2). Taken together, our results show that COREs identified by CREAM share similarities with those identified by ROSE in term of genomic distribution but are associated with stronger differences in gene expression compared to individual CREs.

### CREAM identifies COREs bound by master transcription factors

Transcription factors bind to CREs to modulate the expression of cell-type specific gene expression patterns [35, 36]. COREs were previously found to associate with strong transcription factor binding intensity based on ChIP-seq signal [14]. Hence, we assessed transcription factors binding intensities within COREs using the extensive characterization of transcription factor binding profiles performed by the ENCODE project in GM12878 and K562 cells [30]. We find that more than 25% of transcription factors show binding intensity significantly higher over CREAM-identified COREs compared to individual CREs in both GM12878 and K562 cell lines (FC > 2, FDR < 0.001) (Fig. 3A). In contrast, less than 15% of all transcription factors bind with higher intensity in ROSE-identified COREs compared to individual CREs in both GM12878 and K562 cell lines (FC > 2, FDR < 0.001; Fig. 3A).

**Figure 3.**
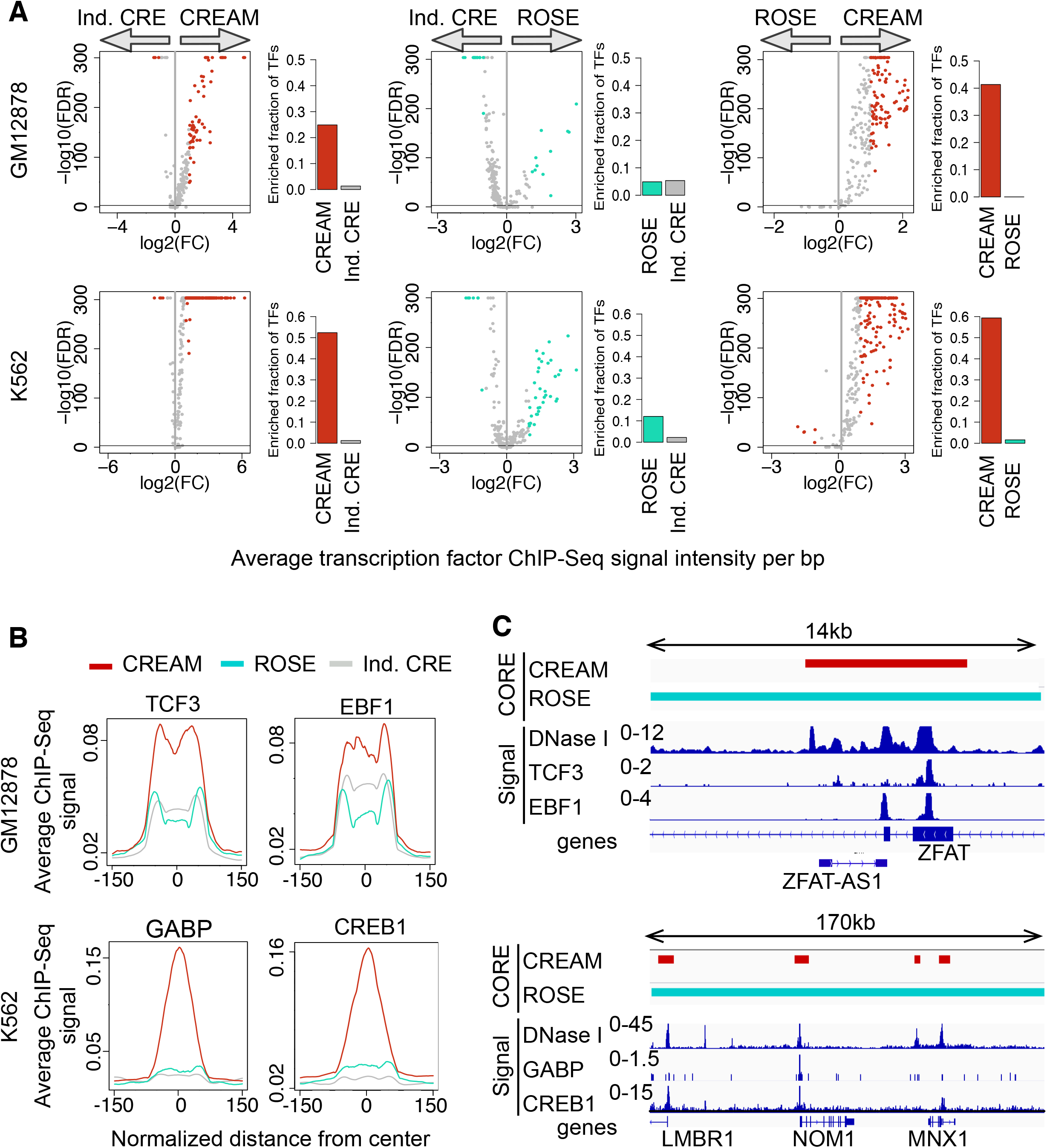
Transcription factor binding enrichment in individual CREs and COREs from GM12878 and K562 cell lines. **A)** Enrichment of transcription factor binding intensity in COREs identified by CREAM or ROSE and individual CREs. Volcano plots represent -log_10_(FDR) versus log_2_(fold change [FC]) in ChIP-seq signal intensities (each dot is one transcription factor) (colored: significant fold change, grey: insignificant fold change). The barplots show how many transcription factors (TFs) have higher signal intensity in COREs or individual CREs (FDR<0.001 and log_2_(FC)>1). Fold change (FC) is defined as the ratio between the average signal per base pair in COREs versus individual CREs or CREAM COREs versus ROSE COREs. **B**) Normalized ChIP-seq signal intensity at COREs and individual CREs for TCF3 and EBF1 as examples of master transcription factors in GM12878 [31] and for GABP and CREB1 as examples of master transcription factors in K562 cells [32–34]. **C**) Examples of genomic regions with COREs identified by both CREAM and ROSE (with different coverage) occupied by transcription factors presented in B.

Difference in transcription factor binding intensity at CREAM versus ROSE-identified COREs is showcased by the master transcription factors TCF3 and EBF1 [31] in GM12878 cells and GABP and CREB1 [32–34] in K562 cells. Indeed, over a 2 fold difference in binding intensity of TCF3 and EBF1 is observed for CREAM-identified COREs compared to individual CREs in GM12878 cell (Fig. 3B). This is exemplified over the CORE proximal to the ZFAT gene in GM12878 cells (Fig. 3C). Similarly, over a 3 fold difference in GABP and CREB1 binding intensity is observed over COREs compared to individual CREs in K562 cells (Fig. 3B) and exemplified at the 7q36 locus harboring a series of COREs bound strongly by GABP and CREB1 in K562 cells (Fig. 3C).

The binding intensity of transcription factors over COREs was calculated as the average ChIP-seq signal within each CORE. We assessed if the difference between enrichment of transcription factor binding intensity within CREAM- and ROSE-identified COREs is not merely due to the difference in their burden of CRE-free gaps. We calculated the transcription factor binding intensity excluding the CRE-free gaps within ROSE-identified COREs (Supplementary Fig. 3). More than 25% of the transcription factors have significantly higher binding intensity within CREAM-identified COREs compared to the signal over the CREs in COREs identified by ROSE (FC > 2, FDR < 0.001).

### CREAM-identified COREs are proximal to essential genes

COREs are reported to lie in proximity to genes essential for self-renewal and pluripotency of stem cells, respectively [37]. A CRISPR/Cas9 gene essentiality screen was recently reported in K562 cells by *Wang et al. (2015)* [38]. Merging these genomic screening data with CORE identification from K562 cells reveals a significant enrichment of gene essential for growth proximal to CREAM-identified COREs (FDR<1e-3 using permutation test; Fig. 4A, B). For instance, the BCR gene is the most essential gene in proximity to a CREAM-identified CORE in K562, a feature of relevance to the expression of the oncogenic BCR-ABL gene fusion reported in Chronic Myelogenous Leukemia [39].

**Figure 4.**
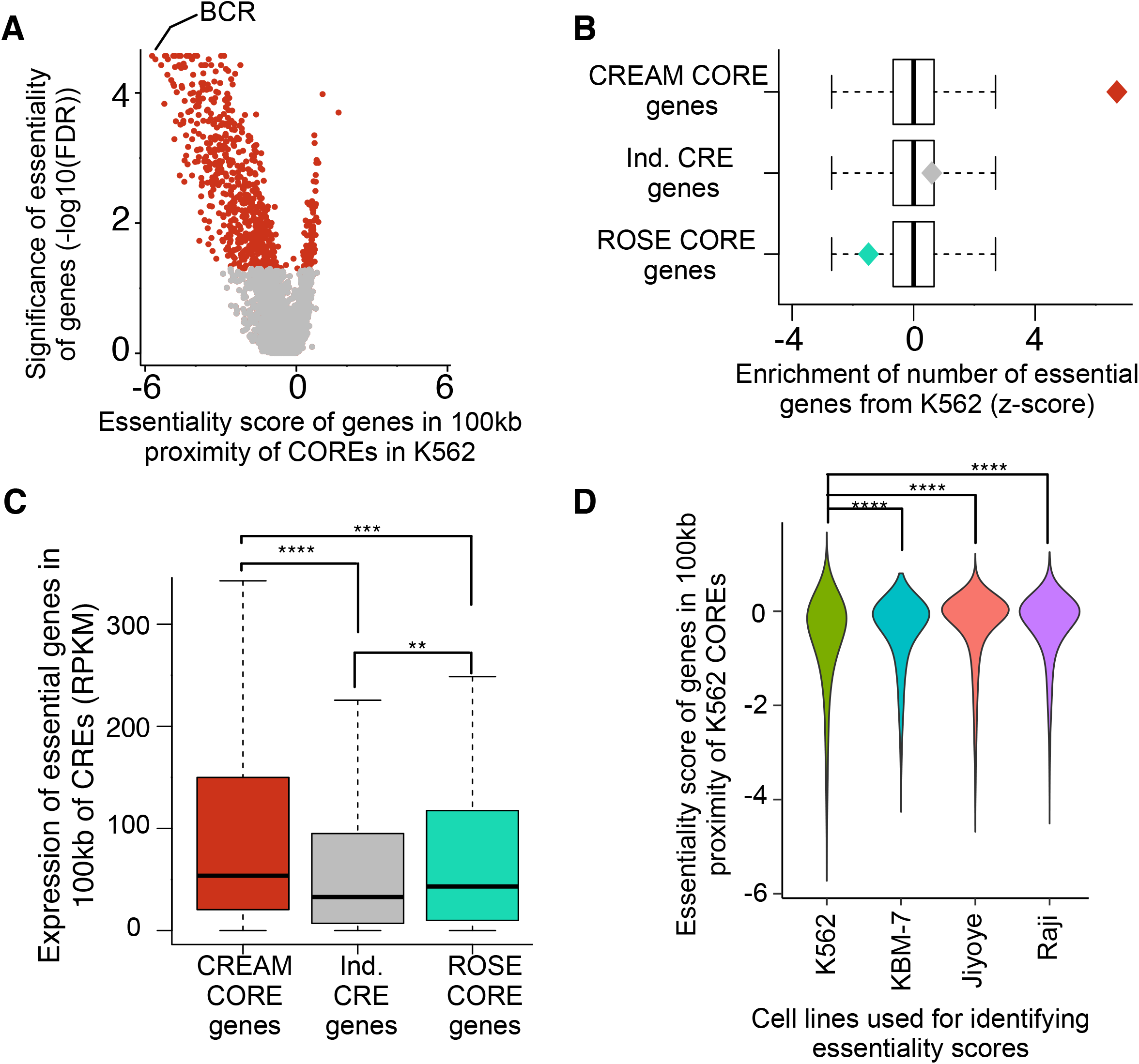
Essentiality of genes in proximity of COREs in K562 cell lines. **(A)** Volcano plot of significance (FDR) and effect size (essentiality score) of genes in proximity of CREAM-identified COREs in K562 cell line (red: significant fold change, grey: insignificant fold change). **(B)** Enrichment based on the number of essential genes among the genes associated to individual CREs and COREs in K562 cell line. (**C**) Transcription level of essential genes associated with individual CREs and COREs. **(D)** Comparing essentiality score of genes reported in K562, KBM-7, Jiyoye, and Raji cell line proximal (100kb) to COREs identified by CREAM in K562.

In contrast, genes proximal to individual CREs are not enriched with essential genes (CRE: FDR=0.26; Fig. 4B), while ROSE-identified COREs are negatively enriched with essential genes (FDR=0.92; Fig. 4B). Expression of genes essential for growth in K562 proximal to CREAM-identified COREs is also significantly higher than the expression of essential genes associated with individual CREs or ROSE-identified COREs (FDR < 0.001; Fig. 4C). Extending our analysis to essentiality scores from other cell lines tested by *Wang et al. (2015)* [38], We show that the essentiality score of genes proximal to K562 CREAM-identified COREs is significantly less in KBM-7, Jiyoye, and Raja cell lines compared to K562 cells (FDR<0.001; Fig. 4D). This supports the cell type-specific nature of COREs and their association with essential genes and argues in favor of COREs accounting for a regulatory potential greater than individual CREs.

### Generalizability of CORE identification across cell and tissue types

ROSE-identified COREs were reported to discriminate cell types [13]. Extended this analysis to CREAM-identified COREs across the DNase-seq defined CREs from 102 cell lines available through the ENCODE project [30] reveals between 1,022 and 7,597 COREs per cell line (Fig. 5A). The number of COREs correlates with the total number of CREs identified in each cell line (Fig. 5B) while fraction of CREs called within COREs is independent of the number of individual CREs (Fig. 5C). However, the average width of COREs across cell lines shows low correlation with the total number of CREs (Spearman correlation ρ<0.25; Fig. 5D), supporting the specificity of CORE widths with respect to each biological sample irrespective of the total number of CREs.

**Figure 5.**
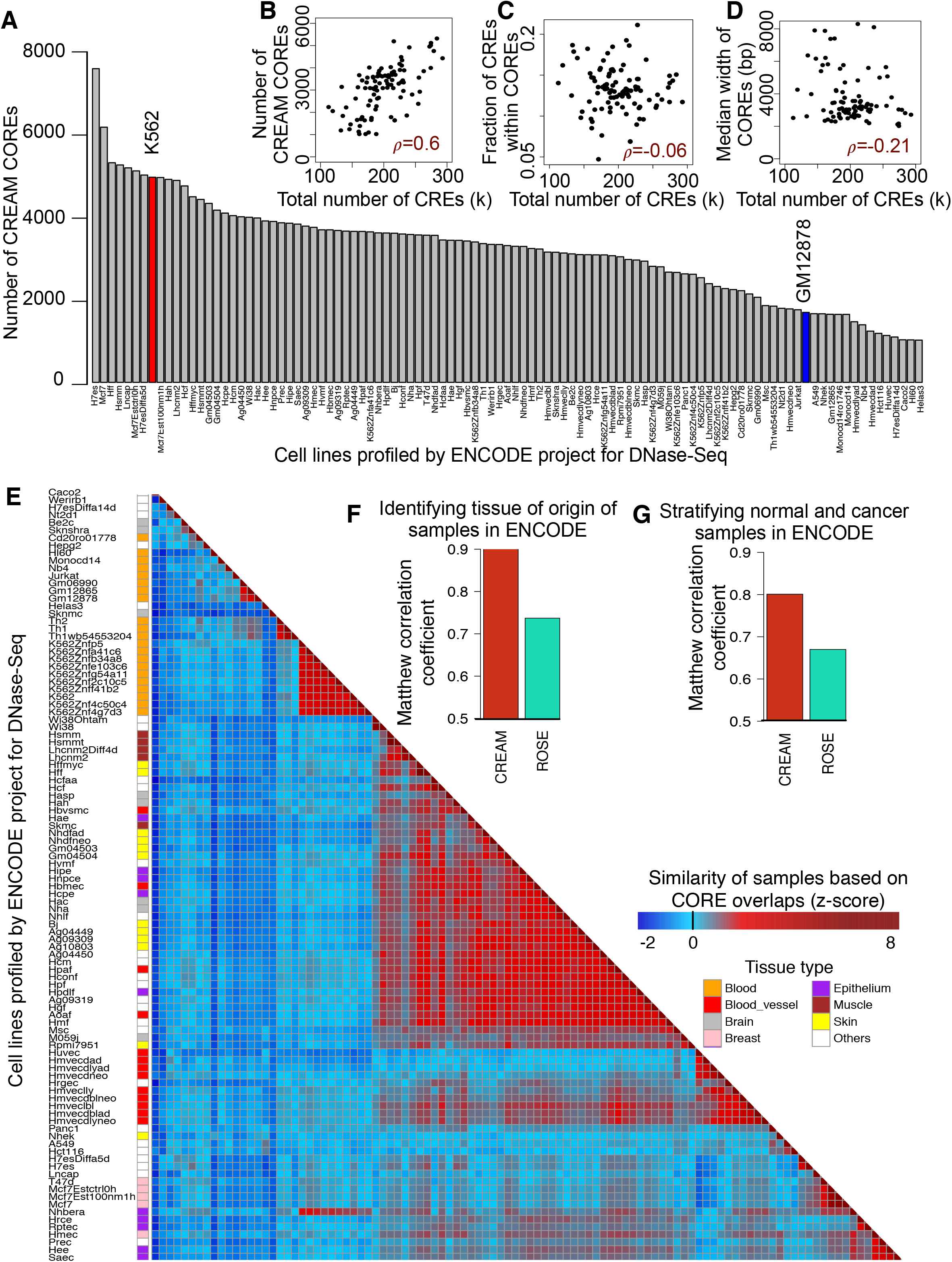
Specificity of COREs to the phenotype of the cell lines in ENCODE. **(A)** Number of COREs identified by CREAM for each cell or tissue previously profiled by the ENCODE project for DNase-seq**. (B)** Positive correlation in the number of COREs with the number of individual CRE per sample. **(C)** Independence in the fraction of CREs called within COREs versus the number of individual CRE per sample. (**D**) Relation between median width of COREs and the number of individual CRE per sample. (**E**) Heatmap of similarities based on CREAM-identified COREs across the ENCODE project cells or tissue samples based on Jaccard index of overlap in COREs. (**F**) Matthew correlation coefficient (MCC) for classification of ENCODE project samples as normal or cancerous using CREAM-versus ROSE-identified COREs. (**G**) Matthew correlation coefficient for classification of ENCODE project samples based on their tissue of origin using CREAM-versus ROSE-identified COREs.

To test whether COREs can discriminate cells with respect to their tissue or origin and their malignant status, we clustered the ENCODE cell lines based on their CREAM- and ROSE-identified COREs. Predicting each cell line based on its nearest neighbor, we could classify tissue of origin with high accuracy using CREAM-identified COREs (Matthew correlation coefficient [MCC] of 0.90 for tissues with ≥ 5 cell lines)(Fig 5E, F). However, ROSE-identified COREs yielded substantially lower predictive value (MCC of 0.74; Fig. 5F). Similarly, CREAM-identified COREs were more discriminative of non-malignant versus malignant cell lines than ROSE-identified COREs (MCC of 0.80 versus 0.67 for CREAM and ROSE, respectively; Fig. 5G).

### Subtyping of tumor samples based on CREAM-identified COREs

We further investigated the utility of CORES to stratify primary tumor samples. We specifically used the H3K27ac ChIP-seq profiles (available in [23]) from eleven TMPRSS2:ERG fusion positive (T2E) and eight fusion negative (non-T2E) primary prostate tumors to identify their COREs based on CREAM and ROSE. Clustering of these prostate tumors samples based on CREAM-identified COREs stratified T2E apart from non-T2E samples (MCC=0.89; Fig. 6A, C), while ROSE COREs were less predictive of tumor samples subtypes (MCC=0.6, Fig. 6B, C). The top predictive COREs discriminating T2E from non-T2E prostate tumor samples map to the ERG genes (Fig. 6D) in agreement with previous report [23], and the SMOC2 gene.

**Figure 6.**
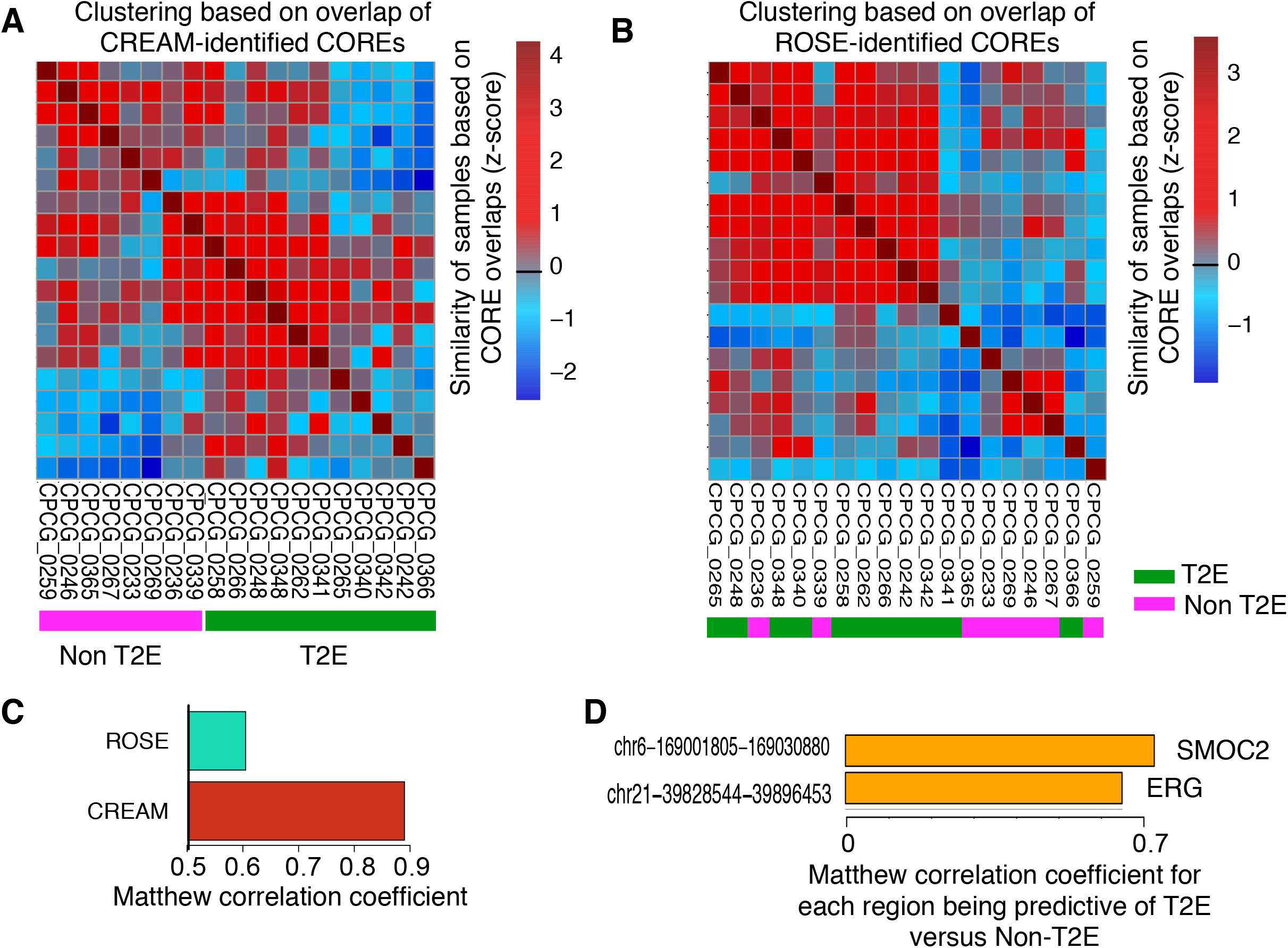
Primary prostate tumor subtyping using COREs. **(A)** Heatmap of similarities across the T2E and non-T2E prostate cancer samples [23] based on Jaccard index of overlap of COREs identified by CREAM on the H3K27ac ChIP-seq in each sample. **(B)** Same as A but based on COREs identified by ROSE. (**C**) Matthew correlation coefficient (MCC) for classification of prostate cancer samples as T2E or non-T2E using CREAM-versus ROSE-identified COREs. (**D**) Top 2 COREs identified by CREAM discriminating T2E from non-T2E primary prostate tumor subtypes.

### Enrichment of COREs at topologically associated domain boundaries

The genome is partitioned into different compartments subdivided into TADs that discriminate active from repressive domains [40]. Here, we integrated the distribution of COREs across the genome with topologically associated domains (TADs), based on publicly available Hi-C data in GM12878 and K562 cell lines [41]. Our analysis reveals higher enrichment of COREs rather than individual CREs at TAD boundaries (Figs. 7A and B). We also report this finding in four other cell lines with available Hi-C data, namely HeLa, HMEC, HUVEC, and NHEK cells [41] (Supplementary Fig. 4). This preferential enrichment of COREs as opposed to individual CREs is not due to difference in size of individual CREs versus COREs (permutation test FDR<0.001, Supplementary Fig. 5) and suggest that TAD boundaries are rich in cis-regulatory elements clustered near each other.

**Figure 7.**
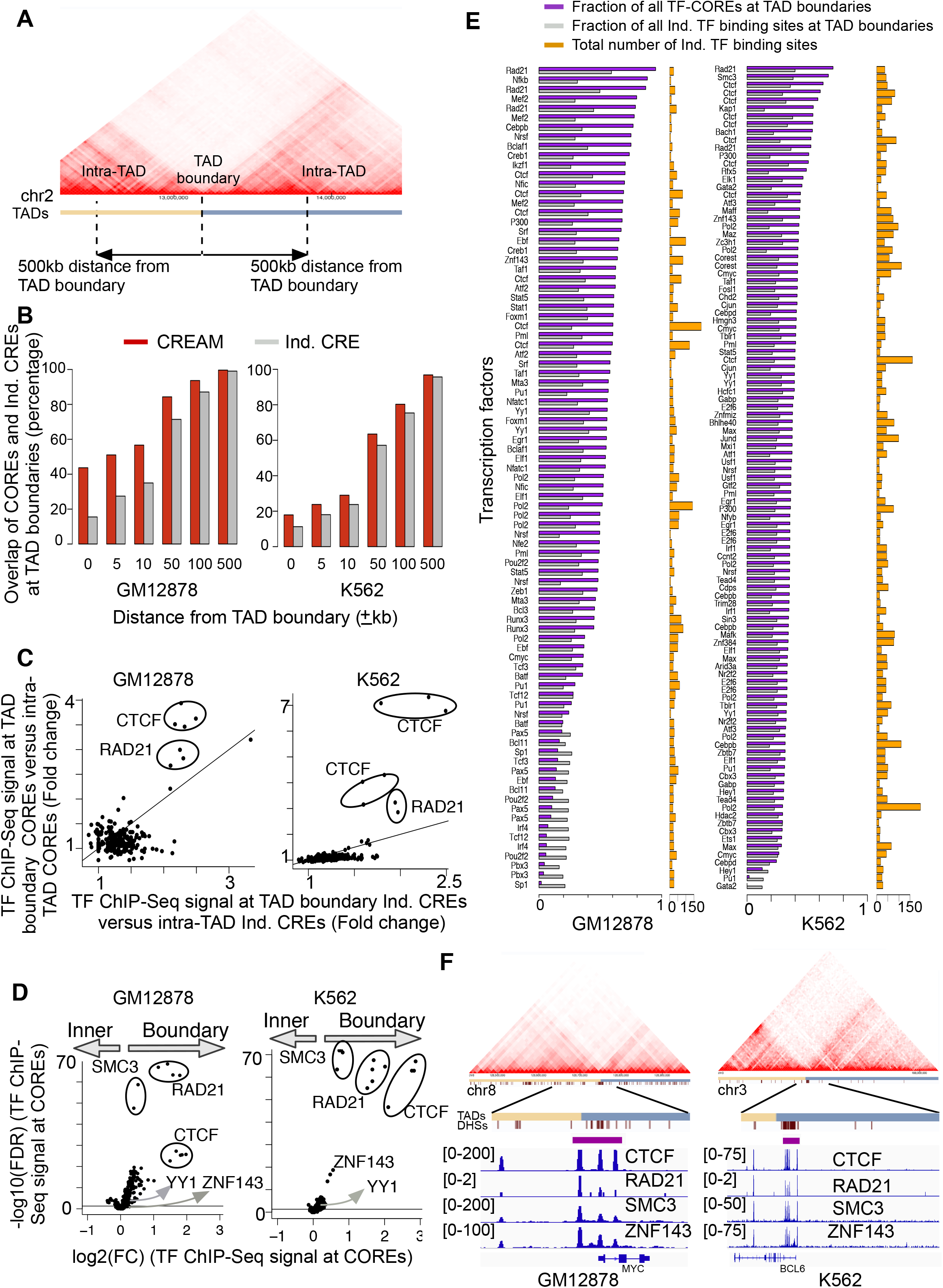
Arrangement of COREs and individual CREs with respect to TAD boundaries. **(A)** Schematic representation of a TAD boundaries and intra-TAD regions (25kb Hi-C resolution). **(B)** Comparison of fraction of COREs and individual CREs from DNAse-seq that lie at TAD boundaries with increasing distance from TAD boundary cutoffs in GM12878 and K562 cell lines. **(C)** Enrichment of transcription factor (TF) binding intensities within COREs versus individual CREs at TAD boundaries (±10kb) versus intra-TADs in GM12878 or K562 cell lines. (**D**) Enrichment of transcription factor binding intensity in TAD-boundary COREs versus intra-TAD COREs 10kb proximity of TAD boundaries (±10kb) versus intra-TAD domains. **(E)** Fraction of transcription factor COREs (TF-COREs: purple) and individual transcription factor binding sites (grey) at TAD boundaries (∓10kb). The total number of individual binding sites for each transcription factor in GM12878 and K562 cell lines is also reported (orange). **(F)** Examples of TF-COREs for CTCF, RAD21, SMC3, and ZNF143 at the TAD boundary for the MYC and BCL6 genes (10kb Hi-C resolution).

CTCF, cohesin (RAD21, SMC1 and SMC3), ZNF143 and the YY1 transcription factors were previously reported to preferentially bind chromatin at TAD boundaries delineating anchors of chromatin interactions [9,41–43]. We therefore assessed if these transcription factors were enrichment within COREs at TAD boundaries based on their ChIP-Seq signal intensity. CTCF and RAD21 were preferential enrichment within COREs rather than individual CREs restricted to TAD boundaries in both GM12878 and K562 cell lines (FC>1.5 for both COREs and Ind. CREs; FC at COREs more than 1.5 times the FC at Ind. CREs; Fig. 7C). No enrichment over COREs at TAD-boundaries was seen for ZNF143 and YY1, or any of the 82 and 94 additional transcription factors with ChIP-seq data in GM12878 and K562 cells, respectively. Together, this argues that CTCF and cohesin behave differently from all other transcription factors at TAD-boundaries, mapping to COREs as opposed to individual CREs. Furthermore, we show that CTCF and cohesin bind at TAD-boundary COREs with higher intensity than at intra-TAD COREs in both GM12878 and K562 cell lines (FC>2, FDR<0.001 for CTCF and RAD21, FC>1.7, FDR<0.001 for SMC3 in GM12878 and K562 respectively; Fig. 7D). ZNF143 also preferentially occupied TAD-boundary COREs as opposed to intra-TAD COREs but only in K562 cells (FC=1.42, FDR < 0.001) (Fig. 7D). We observed lesser differences in the binding intensity of YY1 at TAD-boundary COREs versus intra-TAD COREs in either GM12878 and K562 cell lines (FC<1.25 in both cell lines; Fig. 7D). Extending this analysis to the remaining ChIP-seq data for transcription factors in GM12878 and K562 cell lines [30], revealed 69% and 35% of transcription factors with increased binding intensity at TAD-boundary COREs versus intra-TAD COREs but with low effect size in GM12878 and K562 cell lines, respectively (FC>1, FDR<0.001; Fig. 7D).

The enrichment of CTCF and the cohesin complex within COREs at TAD boundaries led us to assess if they were themselves forming COREs, i.e. present as a cluster of binding sites at TAD boundaries. Using CREAM on the 86 and 98 ChIP-seq data from GM12878 and K562 cells, respectively, identified 41 and 59 transcription factors in either cell line forming at least 100 COREs (Supplementary Table 1). Comparing the distribution of transcription factor-COREs (TF-COREs) at TAD boundaries versus intra-TADs revealed more than 50% of CTCF, RAD21, SMC3, and ZNF143 TF-COREs at TAD boundaries (Fig 7E), exemplified at the *MYC* and *BCL6* gene loci (Fig. 7F). In contrast, less than 10% of the SP1 and GATA2 TF-COREs mapped to TAD boundaries in GM12878 and K562 cell lines, respectively (Fig. 7E). Taken together, these results suggest that clusters of CTCF and cohesin binding sites are preferentially found at TAD boundaries.

### Computational cost

Computationally efficient methods are required to enable comprehensive CORE identification in large-scale studies, such as generated by the ENCODE project. We found that CREAM was on average 42 times faster than ROSE to identify COREs from DNase-seq (Supplementary Fig. 6). To exemplify the impact of running time in typical analyses, we show that CREAM could process 80 DNase-seq data using eight processing cores and 122 giga bytes of random access memory in less than one hour, instead of 27 hours for ROSE.

## CONCLUSIONS

While the concept that CREs are not all equal is well established, their classification into COREs is recent and warrants the development of improved strategies for their classification. Here, we developed CREAM as an unsupervised machine learning method providing a systematic user biased-free approach for identifying COREs. We show that CREAM identifies COREs with higher transcription factor binding intensity and enriched proximal to genes essential for growth compared to individual CREs. CREAM-identified COREs also classify cells according to their tissue of origin, discriminating normal from cancer cells and stratifying tumor subtypes according to their genotype. In all these aspects, CREAM outperforms the ROSE method [20, 29].

We further assessed the biological underpinning of COREs by comparing their distribution with regards to topologically associated domains (TADs), which are reflective of the three-dimensional organization of the genome [41]. Our results show that COREs are enriched compared to individual CREs at TAD boundaries. These COREs are preferentially bound by a limited number of transcription factors, namely CTCF, cohesin(RAD21, SMC3) and to a lesser extent ZNF143. These transcription factors were all reported to bind at anchors of chromatin interaction at TAD boundaries and regulate their formation [9,41–43].

Using CREAM on over 80 transcription factor ChIP-seq data, we report a range in the fraction of binding sites part of COREs. We further show that CTCF and cohesin TF-COREs predominantly map to TAD-boundaries as opposed to intra-TAD regions. Taken together, these results argue for a diverse relationship between COREs and transcription factors, where clusters of chromatin looping factors preferentially accumulate at TAD boundaries. This discovery argues for a model where TAD boundary formation may rely on clusters of CTCF and cohesin binding events.

While our work does not discriminate whether COREs are simply a collection of individual CREs or whether they consist of CREs synergizing with each other, our results highlights the relevance of classifying CREs into individual elements versus COREs to delineate the biology unique to a given sample in either a normal and disease context.

## METHODS

### CREAM

CREAM uses genome-wide maps of cis-regulatory elements (CREs) in the tissue or cell type of interest, such as those generated from chromatin-based assays including DNase-seq, ATAC-seq or ChIP-seq. CREs can be identified from these profiles by peak calling tools such as MACS [44]. The called individual CREs then will be used as input of CREAM. Hence, CREAM does not need the signal intensity files (bam, fastq) as input. CREAM considers proximity of the CREs within each sample to adjust parameters of inclusion of CREs into a CORE in the following steps (Fig. 1):

#### Step 1: Clustering of individual CREs throughout genome

CREAM initially groups neighboring individual CREs throughout the genome. Each group (or cluster) can have different number of individual CREs. Then it categorizes the clusters based on their included CRE numbers. We defined Order (***O***) for each cluster as its included CRE number. In the next steps, CREAM identifies maximum allowed distance between individual CREs for COREs of a given ***O***.

#### Step 2: Maximum window size identification

We defined maximum window size (***MWS***) as the maximum distance between individual CREs included in a CORE. For each ***O***, CREAM estimates a distribution of window sizes, as the maximum distance between individual CREs in all clusters of that ***O*** within the genome. Afterward, ***MWS*** will be identified as follows

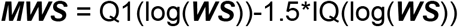

where ***MWS*** is the maximum distance between neighboring individual CREs within a CORE. Q1(log(***WS***)) and IQ(log(***WS***)) are the first quartile and interquartile of distribution of window sizes (Fig. 1).

#### Step 3: Maximum Order identification

After determining ***MWS*** for each Order of COREs, CREAM identifies maximum ***O*** (***O***max) for the given sample. Increasing ***O*** of COREs results in gain of information within the clusters allowing the individual CREs to have further distance from each other. Hence, starting from COREs of ***O***=2, the ***O*** increases up to a plateau at which an increase of ***O*** does not result an increase in ***MWS***. This threshold is considered as maximum ***O*** (***O***max) for COREs within the given sample.

#### Step 4: CORE calling

CREAM starts to identify COREs from ***O***max down to ***O***=2. For each ***O***, it calls clusters with window size less than ***MWS*** as COREs. As a result, many COREs with lower ***O***s are clustered within COREs with higher ***O***s. Therefore, remaining lower ***O*** COREs, for example ***O***=2 or 3, have individual CREs with distance close to ***MWS*** (Fig. 1). These clusters could have been identified as COREs because of the initial distribution of ***MWS*** derived mainly by COREs of the same ***O*** which are clustered in COREs of higher ***O***s. Hence, CREAM eliminate these low ***O*** COREs as follows.

#### Step 5: Minimum Order identification

COREs that contain individual CREs with distance close to ***MWS*** can be identified as COREs due to the high skewness in the initial distribution of ***MWS.*** To avoid reporting these COREs, CREAM filters out the clusters with (***O*** < ***O***min) which does not follow monotonic increase of maximum distance between individual CREs versus **O** (Fig. 1).

### ROSE

ROSE clusters the neighboring individual CREs in a given sample if they have distance less than 12.5kb. It subsequently identifies the signal overlap on the clusters and sorts the identified clusters based on their signal intensity. It then stratifies the clusters based on the inflation point in the sorted clusters and call the clusters with signal intensity higher than the inflation point as super-enhancers (or COREs). This method is comprehensively explained in Whyte *et al.* (2013) [14]. We ran ROSE using the default parameters.

### Genomic overlap of COREs

Bedtools (version 2.23.0) is used to identify unique and shared genomic coverage between CREAM and ROSE-identified COREs.

### Comparison of DNase signal within CORES identified by CREAM and ROSE and single enhancers

Signals (either DNase I hypersensitivity or ChIP-seq) over the identified COREs (or individual CREs) and 1kb flanking regions of them were extracted from the BAM files. Each CORE (or individual CREs) is subsequently binned to 100 binned regions with equal size. Each left and right flanking region is also divided to 100 bins with equal size. Hence, in total 300 bins are obtained for each CORE plus its flanking regions. We then scale the signal in these regions to the library size for the mapped reads. Finally, a Savitzky-Golay filter is applied to remove high frequency noise from the data while preserving the original shape of the data [45, 46]. This filter convolves the signal with a low degree polynomial using least square method [45, 46].

### Association with genes

A gene is considered associated with a CRE or a CORE if found within a ∓100kb window from each other.

### Gene expression comparison

RNA sequencing profile of GM12878 and K562 cells lines, available in ENCODE database [30], are used to identify expression of genes in proximity of individual CREs and COREs. Expression of genes are compared using Wilcoxon signed-rank test.

### Transcription factor binding enrichment

Bedgraph files of ChIP-Seq profiles of transcription factors are overlapped with the identified COREs and individual CREs in GM12878 and K562 using bedtools (version 2.23.0). The resulting signal were summed over all the individual CREs or COREs and then normalized to the total genomic coverage of individual CREs or COREs, respectively. These normalized transcription factor binding intensities are used for comparing TF binding intensity in individual CREs and COREs (Fig. 3). Wilcoxon signed-rank test is used for this comparison.

### Sample similarity

Similarity between two samples in ENCODE is identified based on Jaccard index for the commonality of their identified COREs throughout the genome. Then this Jaccard index is used as the similarity statistics in a 1-nearest-neighbor classification approach. We assess performance of the classification using leave-one-out cross validation. We used Matthew correlation coefficient for performance of the classification model [47]. In this classification scheme, we considered phenotype of the closest sample to an out of pool sample as its phenotype.

### Association with essential genes

Number of genes which are in ∓100kb proximity of COREs and are essential in K562 are identified [38]. This number is then compared with number of essential genes in 10,000 randomly selected (permuted) genes, among the genes included in the essentiality screen. This comparison is used to identify FDR, as number of false discoveries in permutation test, and z-score regarding the significance of enrichment of essential genes among genes in ∓100kb proximity of COREs identified for K562 cell line.

### Enrichment at topologically associated domain

We used HiC profiles generated in GM12878 and K562 cell lines provided by Rao *et al.* (2014) [41]. COREs and individual CREs, from DNase I hypersensitivity profile or ChIP-Seq of transcription factors are then splitted to boundary elements if they were in 10kb proximity of TAD boundaries and intra-TAD if they were further away from the boundaries. We sampled random chromosomal regions with the same size in each chromosome to identify significant of enrichment of COREs of CTCF, RAD21, SMC3, and ZNF143 at boundaries of TADs. We used the 3D genome browser [48] to showcase examples of COREs at boundaries of TADs.

### Research Reproducibility

CREAM is publicly available as an open source R package on the Comprehensive R Archive Network (https://CRAN.R-project.org/package=CREAM).

## List of abbreviations

| Abbreviation | Stand for |
| --- | --- |
| CRE | Cis-Regulatory Element |
| CORE | Cluster Of cis-Regulatory Element |
| CREAM | Clustering of genomic REgions Analysis Method |
| TF | Transcription factor |
| MWS | Maximum Window Size |
| O | Order |
| FC | Fold Change |
| FDR | False Discovery Rate |
| MCC | Matthew Correlation Coefficient |
| TF | Transcription Factor |
| TF-CORE | Cluster of TF binding site |

## DECLARATIONS

### Author contributions

S.A.M.T. developed CREAM. S.A.M.T. and V.K. prepared CREAM R package. S.A.M.T. and P.M. performed the analysis and interpreted the results. S.A.M.T., P.M., B.H-K, and M.L. conceived the design of the study. S.A.M.T., P.M., B.H-K, and M.L. wrote the manuscript. B.H-K and M.L. supervised the study.

### Funding

This study was conducted with the support of the Terry Fox Research Institute, Canadian Cancer Research Society and the Ontario Institute for Cancer Research through funding provided by the Government of Ontario. This work was also supported by the Canadian Institute for Health Research (CIHR: Funding Reference Number 136963 to M.L.) and Princess Margaret Cancer Foundation (M.L. and B.H.K.). We acknowledge the Princess Margaret Bioinformatics group for providing the infrastructure assisting us with analysis presented here. M.L. holds an Investigator Award from the Ontario Institute for Cancer Research, a CIHR New Investigator Award and a Movember Rising Star Award from Prostate Cancer Canada and is proudly funded by the Movember Foundation (grant #RS2014-04). S.A.M.T was supported by Connaught International Scholarships for Doctoral Students. P. M. was supported by the Canadian Institutes of Health Research Scholarship for Doctoral Students. B.H.K is supported by the Gattuso-Slaight Personalized Cancer Medicine Fund at Princess Margaret Cancer Centre and the Canadian Institutes of Health Research.

## Acknowledgment

DNase I sequencing profile of HeLa cell line is used in this research. Henrietta Lacks, and the HeLa cell line that was established from her tumor cells without her knowledge or consent in 1951, have made significant contributions to scientific progress and advances in human health. We are grateful to Henrietta Lacks, now deceased, and to her surviving family members for their contributions to biomedical research. We also acknowledge the ENCODE Consortium and the ENCODE production laboratories that generated the data sets provided by the ENCODE Data Coordination Center used in this manuscript.

## Competing financial interests

The authors declare no competing financial interests.

## REFERENCES

1. Zhou S, Treloar AE, Lupien M. Emergence of the Noncoding Cancer Genome: A Target of Genetic and Epigenetic Alterations. Cancer Discov. 2016;6:1215–29.

2. Kron KJ, Bailey SD, Lupien M. Enhancer alterations in cancer: a source for a cell identity crisis. Genome Med. 2014;6:77.

3. Kellis M, Wold B, Snyder MP, Bernstein BE, Kundaje A, Marinov GK, et al. Defining functional DNA elements in the human genome. Proc Natl Acad Sci U S A. 2014;111:6131–8.

4. Lupien M, Eeckhoute J, Meyer CA, Wang Q, Zhang Y, Li W, et al. FoxA1 translates epigenetic signatures into enhancer-driven lineage-specific transcription. Cell. Elsevier Ltd; 2008;132:958–70.

5. Heintzman ND, Hon GC, Hawkins RD, Kheradpour P, Stark A, Harp LF, et al. Histone modifications at human enhancers reflect global cell-type-specific gene expression. Nature. Nature Publishing Group; 2009;459:108–12.

6. Ernst J, Kheradpour P, Mikkelsen TS, Shoresh N, Ward LD, Epstein CB, et al. Mapping and analysis of chromatin state dynamics in nine human cell types. Nature. Nature Publishing Group; 2011;473:43–9.

7. Heintzman ND, Stuart RK, Hon G, Fu Y, Ching CW, Hawkins RD, et al. Distinct and predictive chromatin signatures of transcriptional promoters and enhancers in the human genome. Nat Genet. Nature Publishing Group; 2007;39:311–8.

8. Hnisz D, Abraham BJ, Lee TI, Lau A, Saint-André V, Sigova AA, et al. Super-enhancers in the control of cell identity and disease. Cell. 2013;155:934–47.

9. Bailey SD, Zhang X, Desai K, Aid M, Corradin O, Cowper-Sal Lari R, et al. ZNF143 provides sequence specificity to secure chromatin interactions at gene promoters. Nat Commun. 2015;2:6186.

10. Thurman RE, Rynes E, Humbert R, Vierstra J, Maurano MT, Haugen E, et al. The accessible chromatin landscape of the human genome. Nature. Nature Publishing Group; 2012;489:75–82.

11. Buenrostro JD, Giresi PG, Zaba LC, Chang HY, Greenleaf WJ. Transposition of native chromatin for fast and sensitive epigenomic profiling of open chromatin, DNA-binding proteins and nucleosome position. Nat Methods. Nature Publishing Group; 2013;1–8.

12. Parker SCJ, Stitzel ML, Taylor DL, Orozco JM, Erdos MR, Akiyama JA, et al. Chromatin stretch enhancer states drive cell-specific gene regulation and harbor human disease risk variants. Proc Natl Acad Sci U S A. 2013;110:17921–6.

13. Hnisz D, Abraham BJ, Lee TI, Lau A, Saint-André V, Sigova AA, et al. Super-enhancers in the control of cell identity and disease. Cell. 2013;155:934–47.

14. Whyte WA, Orlando DA, Hnisz D, Abraham BJ, Lin CY, Kagey MH, et al. Master transcription factors and mediator establish super-enhancers at key cell identity genes. Cell. 2013;153:307–19.

15. Gaulton KJ, Nammo T, Pasquali L, Simon JM, Giresi PG, Fogarty MP, et al. A map of open chromatin in human pancreatic islets. Nature Publishing Group. Nature Publishing Group; 2010;42:255–9.

16. Dowen JM, Fan ZP, Hnisz D, Ren G, Abraham BJ, Zhang LN, et al. Control of cell identity genes occurs in insulated neighborhoods in mammalian chromosomes. Cell. 2014;159:374–87.

17. Vahedi G, Kanno Y, Furumoto Y, Jiang K, Parker SCJ, Erdos MR, et al. Super-enhancers delineate disease-associated regulatory nodes in T cells. Nature. 2015;520:558–62.

18. Corradin O, Cohen AJ, Luppino JM, Bayles IM, Schumacher FR, Scacheri PC. Modeling disease risk through analysis of physical interactions between genetic variants within chromatin regulatory circuitry. Nat Genet. 2016;48:1313–20.

19. Pasquali L, Gaulton KJ, Rodríguez-Seguí SA, Mularoni L, Miguel-Escalada I, Akerman İ, et al. Pancreatic islet enhancer clusters enriched in type 2 diabetes risk-associated variants. Nat Genet. 2014;46:136–43.

20. Lovén J, Hoke HA, Lin CY, Lau A, Orlando DA, Vakoc CR, et al. Selective inhibition of tumor oncogenes by disruption of super-enhancers. Cell. 2013;153:320–34.

21. Northcott PA, Lee C, Zichner T, Stütz AM, Erkek S, Kawauchi D, et al. Enhancer hijacking activates GFI1 family oncogenes in medulloblastoma. Nature. 2014;511:428–34.

22. Chipumuro E, Marco E, Christensen CL, Kwiatkowski N, Zhang T, Hatheway CM, et al. CDK7 inhibition suppresses super-enhancer-linked oncogenic transcription in MYCN-driven cancer. Cell. 2014;159:1126–39.

23. Kron KJ, Murison A, Zhou S, Huang V, Yamaguchi TN, Shiah Y-J, et al. TMPRSS2–ERG fusion co-opts master transcription factors and activates NOTCH signaling in primary prostate cancer. Nat Genet. Nature Publishing Group, a division of Macmillan Publishers Limited. All Rights Reserved.; 2017;49:1336.

24. Moorthy SD, Davidson S, Shchuka VM, Singh G, Malek-Gilani N, Langroudi L, et al. Enhancers and super-enhancers have an equivalent regulatory role in embryonic stem cells through regulation of single or multiple genes. Genome Res. 2017;27:246–58.

25. Hay D, Hughes JR, Babbs C, Davies JOJ, Graham BJ, Hanssen L, et al. Genetic dissection of the α-globin super-enhancer in vivo. Nat Genet. 2016;48:895–903.

26. Shin HY, Willi M, HyunYoo K, Zeng X, Wang C, Metser G, et al. Hierarchy within the mammary STAT5-driven Wap super-enhancer. Nat Genet. 2016;48:904–11.

27. Dukler N, Gulko B, Huang Y-F, Siepel A. Is a super-enhancer greater than the sum of its parts? Nat Genet. 2016;49:2–3.

28. Pott S, Lieb JD. What are super-enhancers? Nat Genet. Nature Publishing Group; 2014;47:ng.3167.

29. Whyte WA, Orlando DA, Hnisz D, Abraham BJ, Lin CY, Kagey MH, et al. Master Transcription Factors and Mediator Establish Super-Enhancers at Key Cell Identity Genes. Cell. Elsevier Inc.; 2013;153:307–19.

30. ENCODE Project Consortium. An integrated encyclopedia of DNA elements in the human genome. Nature. 2012;489:57–74.

31. Somasundaram R, Prasad MAJ, Ungerbäck J, Sigvardsson M. Transcription factor networks in B-cell differentiation link development to acute lymphoid leukemia. Blood. 2015;126:144–52.

32. Erkeland SJ, Valkhof M, Heijmans-Antonissen C, Delwel R, Valk PJM, Hermans MHA, et al. The gene encoding the transcriptional regulator Yin Yang 1 (YY1) is a myeloid transforming gene interfering with neutrophilic differentiation. Blood. 2003;101:1111–7.

33. Yang Z-F, Zhang H, Ma L, Peng C, Chen Y, Wang J, et al. GABP transcription factor is required for development of chronic myelogenous leukemia via its control of PRKD2. Proc Natl Acad Sci U S A. 2013;110:2312–7.

34. Shankar DB, Cheng JC, Kinjo K, Federman N, Moore TB, Gill A, et al. The role of CREB as a proto-oncogene in hematopoiesis and in acute myeloid leukemia. Cancer Cell. 2005;7:351–62.

35. Bulger M, Groudine M. Functional and mechanistic diversity of distal transcription enhancers. Cell. 2011;144:327–39.

36. Maston GA, Evans SK, Green MR. Transcriptional regulatory elements in the human genome. Annu Rev Genomics Hum Genet. 2006;7:29–59.

37. Di Micco R, Fontanals-Cirera B, Low V, Ntziachristos P, Yuen SK, Lovell CD, et al. Control of embryonic stem cell identity by BRD4-dependent transcriptional elongation of super-enhancer-associated pluripotency genes. Cell Rep. 2014;9:234–47.

38. Wang T, Birsoy K, Hughes NW, Krupczak KM, Post Y, Wei JJ, et al. Identification and characterization of essential genes in the human genome. Science. 2015;350:1096–101.

39. Ren R. Mechanisms of BCR--ABL in the pathogenesis of chronic myelogenous leukaemia. Nat Rev Cancer. Nature Publishing Group; 2005;5:172–83.

40. Ea V, Baudement M-O, Lesne A, Forné T. Contribution of Topological Domains and Loop Formation to 3D Chromatin Organization. Genes. 2015;6:734–50.

41. Rao SSP, Huntley MH, Durand NC, Stamenova EK, Bochkov ID, Robinson JT, et al. A 3D map of the human genome at kilobase resolution reveals principles of chromatin looping. Cell. 2014;159:1665–80.

42. Heidari N, Phanstiel DH, He C, Grubert F, Jahanbani F, Kasowski M, et al. Genome-wide map of regulatory interactions in the human genome. Genome Res. 2014;24:1905–17.

43. Weintraub AS, Li CH, Zamudio AV, Sigova AA, Hannett NM, Day DS, et al. YY1 Is a Structural Regulator of Enhancer-Promoter Loops. Cell. 2017;171:1573–88.e28.

44. Feng J, Liu T, Zhang Y. Using MACS to identify peaks from ChIP-Seq data. Curr Protoc Bioinformatics. 2011;Chapter 2:Unit 2.14.

45. Savitzky A, Golay MJE. Smoothing and Differentiation of Data by Simplified Least Squares Procedures. Anal Chem. American Chemical Society; 1964;36:1627–39.

46. Press WH. Numerical Recipes in C: The Art of Scientific Computing. Cambridge University Press; 1992.

47. Safikhani Z, Smirnov P, Freeman M, El-Hachem N, She A, Rene Q, et al. Revisiting inconsistency in large pharmacogenomic studies. F1000Res. 2016;5:2333.

48. Wang Y, Zhang B, Zhang L, An L, Xu J, Li D, et al. The 3D Genome Browser: a web-based browser for visualizing 3D genome organization and long-range chromatin interactions [Internet]. bioRxiv. 2017 [cited 2018 Feb 2]. p. 112268. Available from: https://www.biorxiv.org/content/early/2017/02/27/112268.abstract

