## Supplementary Materials for "CREAM: Clustering of genomic REgions Analysis Method"

**Supplementary Table 1.** Number of TF-COREs identified by CREAM for transcription factors with ChIP-Seq profiles in ENCODE project for GM12878 and K562 and more than 100 TF-COREs.

| GM12878 | | K562 | |
|---|---|---|---|
| Transcription factor | Average number of TF-COREs among replicates | Transcription factor | Average number of TF-COREs among replicates |
| RUNX3 | 1334 | MAFK | 904 |
| CTCF | 1284 | MAFF | 732 |
| ZNF143 | 843 | JUND | 685 |
| EBF1 | 793 | ZNF143 | 673 |
| CMYC | 767 | TEAD4 | 657 |
| RAD21 | 693 | RAD21 | 646 |
| EBF | 677 | CTCF | 568 |
| STAT1 | 593 | PU1 | 544 |
| YY1 | 564 | ELF1 | 514 |
| MTA3 | 531 | MAX | 498 |
| BATF | 428 | HDAC2 | 494 |
| TCF12 | 397 | CBX3 | 481 |
| STAT5 | 388 | CCNT2 | 477 |
| P300 | 385 | BHLHE40 | 470 |
| BCL3 | 384 | HCFC1 | 463 |
| ATF2 | 382 | ZNF384 | 463 |
| PAX5 | 376 | IRF1 | 456 |
| FOXM1 | 375 | SMC3 | 450 |
| PU1 | 371 | TBLR1 | 446 |
| NFIC | 368 | CEBPB | 442 |
| BCLAF1 | 340 | CMYC | 429 |

| | |
|---|---|
| NFATC1 | 339 |
| PML | 321 |
| TCF3 | 295 |
| SP1 | 272 |
| ELF1 | 236 |
| IRF4 | 232 |
| IKZF1 | 189 |
| POU2F2 | 178 |
| BCL11 | 177 |
| TAF1 | 168 |
| POL2 | 165 |
| MEF2 | 162 |
| ZEB1 | 156 |
| PBX3 | 150 |
| NFE2 | 138 |
| CREB1 | 134 |
| SRF | 132 |
| CEBPB | 129 |
| EGR1 | 109 |
| NRSF | 106 |
| NFKB | 103 |

| | |
|---|---|
| NRSF | 423 |
| CDPS | 403 |
| P300 | 378 |
| ZNFMIZ | 376 |
| NR2F2 | 364 |
| ARID3A | 362 |
| EGR1 | 352 |
| RFX5 | 347 |
| ZBTB7 | 338 |
| HEY1 | 321 |
| COREST | 313 |
| PML | 308 |
| ATF1 | 305 |
| MAZ | 276 |
| CJUN | 244 |
| CHD2 | 243 |
| CEBPD | 236 |
| USF1 | 226 |
| MXI1 | 219 |
| ZC3H1 | 218 |
| TRIM28 | 209 |
| E2F6 | 208 |
| NFYB | 205 |
| STAT5 | 203 |
| YY1 | 198 |
| KAP1 | 189 |

| | |
|---|---|
| ELK1 | 187 |
| POL2 | 180 |
| GABP | 156 |
| ATF3 | 156 |
| HMGN3 | 154 |
| GATA2 | 153 |
| BACH1 | 151 |
| TAF1 | 148 |
| SIN3 | 145 |
| ETS1 | 139 |
| GTF2 | 129 |
| FOSL1 | 110 |

**Supplementary Figure 1**

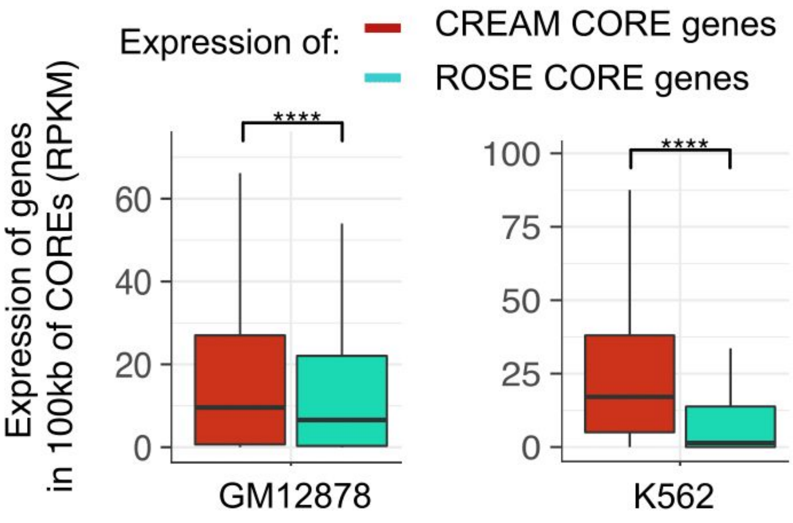

**Supplementary Figure 2**

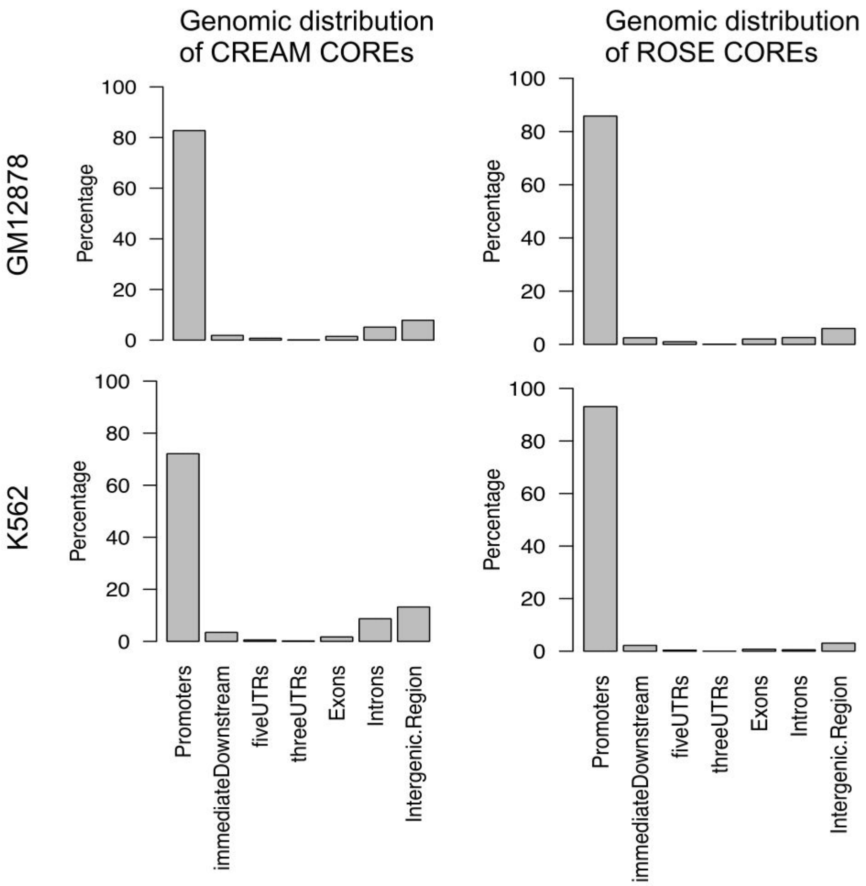

**Supplementary Figure 3**

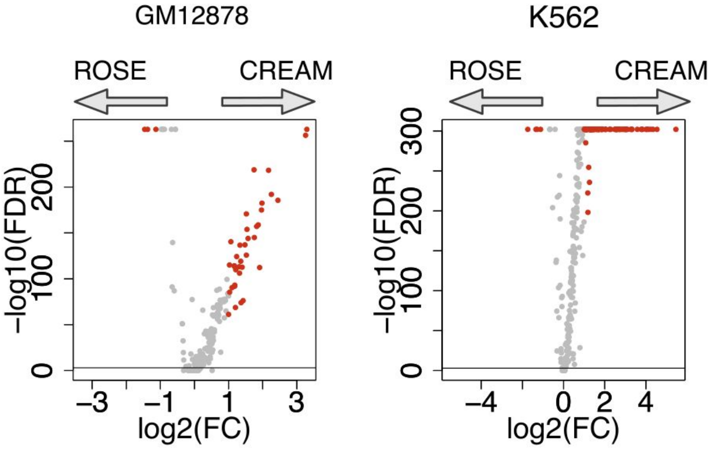


Average transcription factor ChIP-Seq signal intensity per bp

**Supplementary Figure 4**

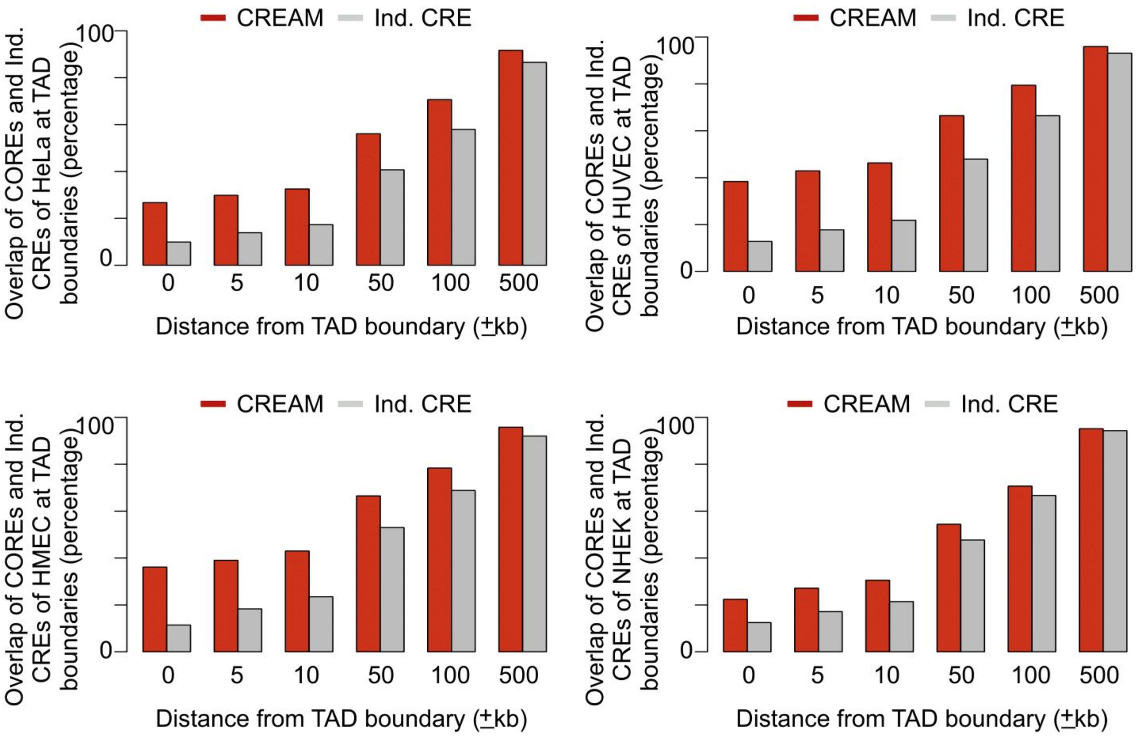

**Supplementary Figure 5**

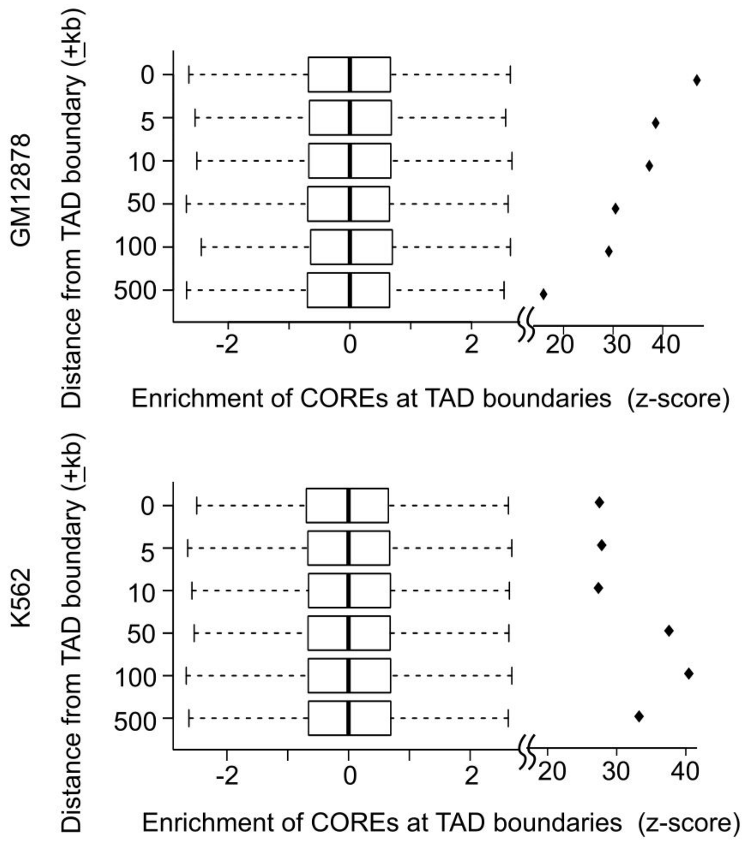

**Supplementary Figure 6**

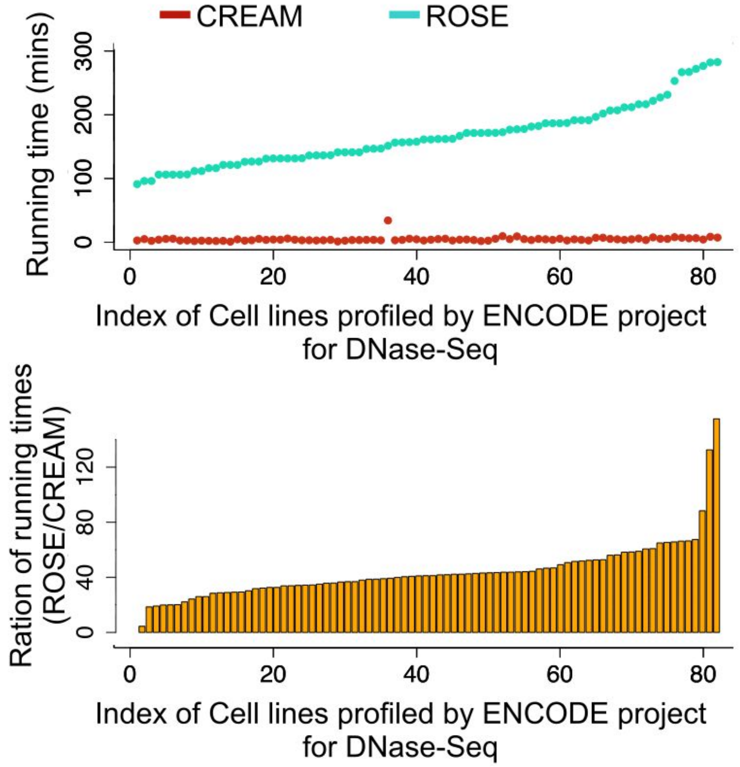
